# Phasic dopamine enhances the distinct decoding and perceived salience of stimuli

**DOI:** 10.1101/771162

**Authors:** Lars-Lennart Oettl, Max Scheller, Sebastian Wieland, Franziska Haag, David Wolf, Cathrin Loeb, Namasivayam Ravi, Daniel Durstewitz, Roman Shusterman, Eleonora Russo, Wolfgang Kelsch

## Abstract

Subjects learn to assign value to stimuli that predict outcomes. Novelty, rewards or punishment evoke reinforcing phasic dopamine release from midbrain neurons to ventral striatum that mediates expected value and salience of stimuli in humans and animals. It is however not clear whether phasic dopamine release is sufficient to form distinct engrams that encode salient stimuli within these circuits. We addressed this question in awake mice. Evoked phasic dopamine induced plasticity selectively to the population encoding of coincidently presented stimuli and increased their distinctness from other stimuli. Phasic dopamine thereby enhanced the decoding of previously paired stimuli and increased their perceived salience. This dopamine-induced plasticity mimicked population coding dynamics of conditioned stimuli during reinforcement learning. These findings provide a network coding mechanism of how dopaminergic learning signals promote value assignment to stimulus representations.

## Introduction

We experience a plethora of sensory stimuli in our environment. Only some of these cues, however, turn out to be predictive of relevant situations. Initially neutral stimuli become more attended and elicit reactions when associated with rewarding or aversive events, thereby rendering the stimuli more salient over others. These processes shape most learnt behaviors ranging from social relations (1–3) to major psychiatric disorders with learnt cues causing drug seeking (4–8) or aberrant salience states in the development of psychosis (9, 10).

Dopaminergic midbrain neurons fire brief bursts in response to rewards and provide phasic dopamine release (pDA) as a reinforcement signal to cortico-striatal circuits (11–15). Cortico-striatal circuits encode different features of stimuli (16, 17). In these circuits, value and identity information are dynamically encoded by populations of neurons with excitatory and inhibitory responses that evolve over time in response to the stimulus (18–20). While some of the primary sensory areas mainly code for the identity of the stimulus (21–24), the expected value of a stimulus has been extensively shown to be represented in the ventral striatum in human imaging studies and recordings from many other species (25–28). In particular, neuronal activity in the ventral striatum is modified when the subject learns to predict future rewards from sensory cues (27–29).

Mesolimbic dopamine in the ventral striatum is critical to reinforcement learning (30, 31) and the incentive salience hypothesis of dopamine function in reward suggests that dopaminergic neurotransmission in the ventral striatum modulates the attribution of salience to conditioned cues that anticipate reward (25). While the behavioral sufficiency of dopamine to render stimuli more salient is well established (32–34), it is not clear in how far pDA modifies directly the neuronal population encoding of stimuli in ventral striatal networks. We therefore tested in awake animals whether pDA release itself is sufficient to induce plasticity to the population encoding of initially neutral stimuli to generate distinct stimulus representations with increased perceived salience. These findings may contribute to understand how reinforcement signals modify encoding in neuronal networks for value assignment to stimuli.

## Results

### Value assignment to ventral striatal stimulus responses during reinforcement learning

We aimed to understand how pDA modifies stimulus-triggered representations in the ventral striatum. The ventral striatum is composed of the olfactory tubercle (OTu) and nucleus accumbens (Fig. 1a). The OTu provides the main direct access of olfactory information to the limbic reward system (35) and shares functions in reinforcement and salience with nucleus accumbens (34–36). In the first instance, we examined how neuronal representations are modified during a reversal learning go/no-go task. Specifically, we asked whether task-learning affects the population response to the rewarded odorant and whether the neuronal representations of the rewarded and non-rewarded odorant become more distinct with learning. We focused on the odor-cue triggered responses occurring between stimulus onset and reward delivery.

**Figure 1.**
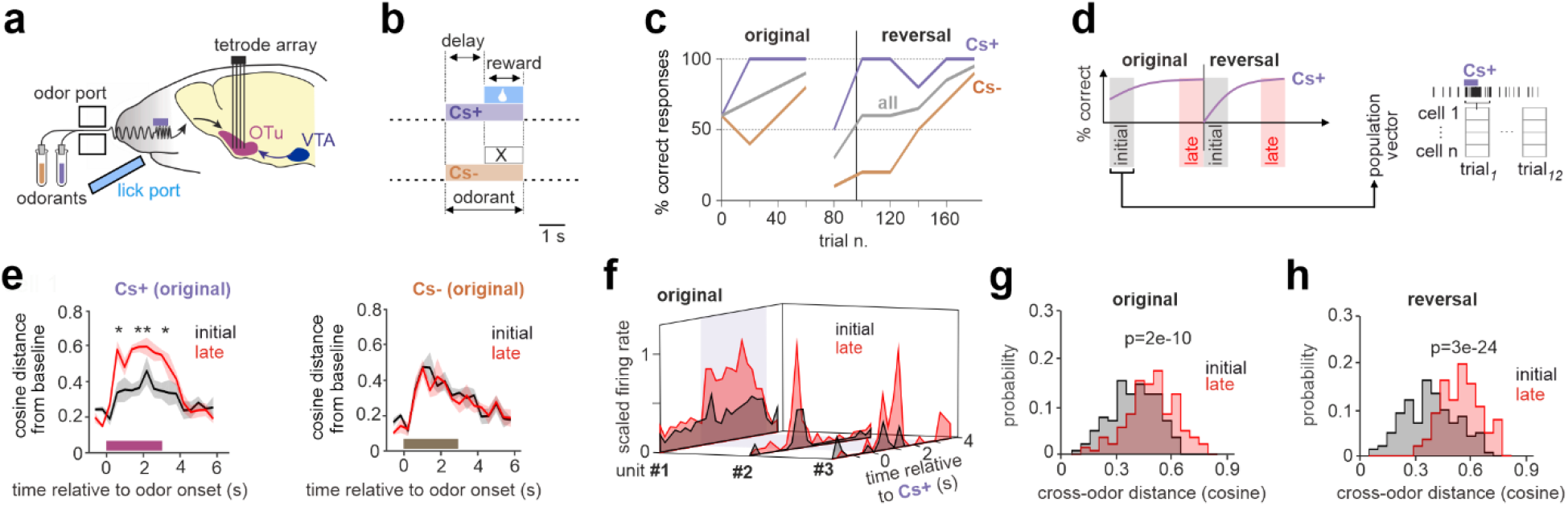
Value assignment to sensory stimuli in the ventral striatum during reversal learning. (a) To assess how stimulus-triggered neuronal responses are modified during reversal learning, we performed recordings in head fixed mice with a newly designed tetrode array, placed in the olfactory tubercle (OTu) of the ventral striatum. Odorants were delivered through an olfactometer and licking for water rewards was detected by an IR beam breaker sensor. (b) Mice learnt to perform a go/no-go task with a water drop being delivered when licking upon Cs+ presentation. No water drop was obtained when licking during Cs-presentation. Odorants were presented for 3 s and only licking after >1.5 s after odorant onset evoked release of the water drop. (c) In the same session, mice learnt to perform to criterion (80% combined hits and correct rejections within 40 consecutive trials), then the other odorant was rewarded, and animals again learnt to criterion. (d) To reveal changes after learning in the population coding in response to Cs+ and Cs-, we divided the recording session into an ‘initial’ phase (first 12 Cs+ and Cs-trials) and a ‘late’ phase (last twelve Cs+ and Cs-trials). For each trial of these phases, we collected network activity in a population vector. (e) Mean temporal evolution ± standard error of the cosine distance of the population vector from baseline for the rewarded and non-rewarded odorant for the ‘initial’ (black) and ‘late’ (red) phase of the session. Significant changes between the ‘initial’ and ‘late’ phase were assessed with a two-tailed t-test corrected for multiple comparisons across bins. During learning the population vector for Cs+, but not for Cs-, significantly increased its distance from baseline. (f) Examples of 3 pSPNs with responses to Cs+ in the ‘initial’ phase of the session (average of the first 12 Cs+ trials) and ‘late’ after reaching criterion (last twelve Cs+ trials). The responses were normalized to the peak response (250 ms binning) of each unit. (g) Distribution of distances between the trial-specific neuronal representations of Cs+ and Cs-within all trials of the ‘initial’ phase (black) and all trials of the ‘late’ phase (red). Distances were computed both with a cosine metrics. Significance was assessed with a two-tailed t-test. During learning the population vectors relative Cs+ and Cs-diverged. (g) same as (h) but computed on trials following the reversal of odorant-reward contingency.

We recorded from the OTu (Fig. 1a, SI Fig. S1a-b) with a newly designed custom-built chronically implanted tetrode array that allowed for recordings with up to 128 channels. The array, including connectors, had a total weight of less than 2 grams and was of approximately 5 mm in height (SI Fig. S1c-d). Recorded mice performed a go/no-go task in which they learnt to lick during the rewarded odorant and not to lick during the non-rewarded one. Once the animal reached criterion (80% correct responses over 40 trials), reward contingencies were reversed within the same session and the new association learnt (Fig. 1b-c). Phasic DA-release occurs prominently during reward retrieval (12, 15); we therefore structured the task so that the odorants were presented for 3 s and the reward could be retrieved after 1.5 s from stimulus onset (Fig. 1b).

From a total of 75 recorded single units, 28 units had baseline firing rates below 5 Hz and were further analyzed as putative striatal projection neurons (pSPN) (SI Fig. S1e). These units displayed features compatible with those previously reported for SPNs throughout striatum (37). The majority of pSPNs did maximally respond during the first second after odorant onset (SI Fig. S1f). To study the effect of learning on odor encoding we compared the neuronal responses of pSPNs from the first 12 trials of the session, when animals were still unsure about the association of odor and reward, to the last 12 trials, when animals had reached criterion. The stimulus responses of individual pSPNs were both excitatory and inhibitory (SI Fig. S1g). To better capture the dynamics, we addressed this question at the population level by assembling sets of pSPNs across experiments into high dimensional population vectors (Fig. 1d). The advantage of using a population approach is that it allows for accounting even for small changes in rate, when coherent, by collecting them throughout the whole population. Further, when neuronal responses are both excitatory and inhibitory, like in the present sample, averaging across neurons may cancel out the network response. The population analysis was designed to account for both of these problems. Population vectors of units were concatenated across animals separately for Cs+ and Cs-, thereby maintaining the trial temporal evolution. To control for the possibility that the specific concatenation of trials across sessions influenced our results, we repeated all main analyses with 300 random permutations of cross-session trial matching (SI Fig. S2). To quantify the deflection of the network activity from baseline following odor stimulation, we computed the distance of the population vector from the average-baseline-vector during the whole trial (see Methods). Distances were computed both with a cosine and a Euclidean metric to account for differences in angle and rate of the two neural activity vectors. The analysis revealed that, while no changes occurred in response to the non-rewarded odorant (Fig. 1e), the neural representation of the rewarded odor became more distinct from baseline while the animals learnt to assign value to the stimulus (Fig. 1e) (see also SI Figure 2a). Within the sample supra-threshold excitatory responses to Cs+ also increased during the reversal learning (Fig. 1f). As predicted, the population vectors of the trials paired to the rewarded odorant and those paired to the non-rewarded odorant were more dissimilar after learning (Fig. 1g) (see also SI Figure S2d-e). During the reversal phase, a similar effect was observed (Fig. 1h) (see also SI Figure S2d,f). The re-learning of the odorant-reward association increased the odor response to the now rewarded odorant (SI Fig. S2b-c). Only in the reversal learning, the response to the non-rewarded odorant decreased (SI Fig. S2b-c).

**Figure 2.**
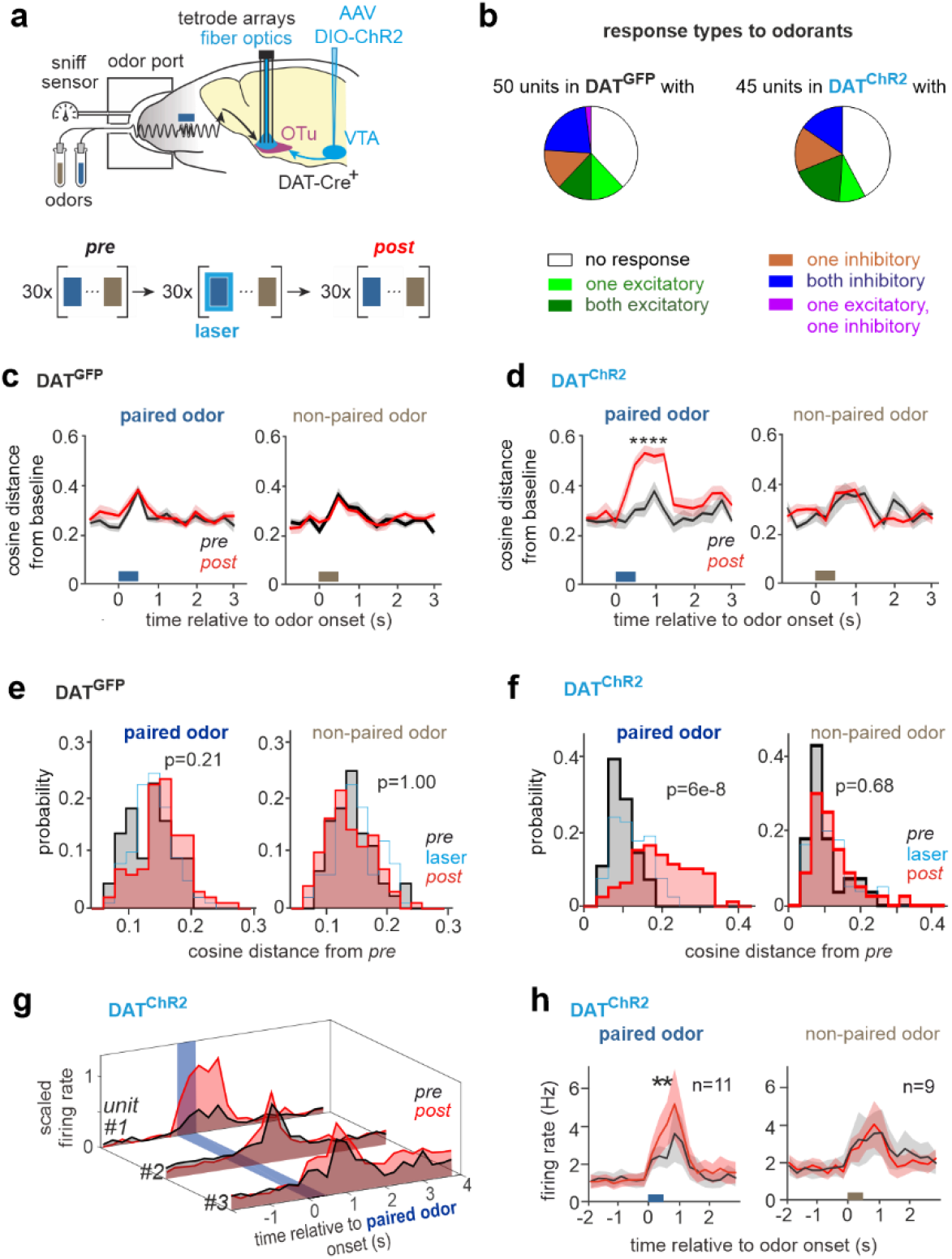
Phasic DA modifies the striatal population encoding selectively of the paired odorant. (a) ChR2 was conditionally expressed by AAV delivery to the VTA of DAT:Cre mice (DAT^ChR2^). Recordings were performed with chronic tetrode arrays and fiber optics targeting bilaterally the OTu. Mice were head fixed and odorants were applied through a mask that also allowed for monitoring of the sniff cycle. (Bottom) Two odorants were applied for 0.5 s in pseudorandomized order every 10 s. During the ‘pairing’ phase, one odorant was consistently paired with laser light delivered to the OTu to evoke DA release. Sham laser light trains (40 Hz for 300 ms) were applied the animals’ heads during all other odorant presentations including the ‘non-paired’ odorant. (b) In our sample, we obtained 50 stable pSPNs in mice that expressed GFP in DA neurons (DAT^GFP^) and 45 pSPNs in DAT^ChR2^ mice. The pie chart visualizes the relative occurrence of either inhibitory or excitatory responses to one or both of the odorants. (c-d) Mean temporal evolution ± standard error of the cosine distance of the population vector from baseline for the two odorants during the ‘pre’ and ‘post’ phases for (c) DAT^GFP^ and (d) DAT^ChR2^ mice. Significant changes between the two phases were assessed with a two-tailed t-test corrected for multiple comparisons across bins. Phasic DA-paring increased the distance of the odor response from baseline exclusively for the paired odorant and in DAT^ChR2^ mice. (e-f) Comparison between the distribution of cosine distances between response vectors within the ‘pre’ phase (black) and between the ‘pre’ and ‘post’ phase (red), for (e) DAT^GFP^ and (f) DAT^ChR2^ mice. Significance established with a three-way ANOVA (factors: cohort, phase, and odor). Interaction effect: F(1,498)=8.0; p=0.005. Post-hoc tests (Tukey’s correction) are reported on the plots that show that only DAT^ChR2^ mice show a significant change in odor response to the paired odorant after pDA paring. (g) Example of peri-stimulus histograms (PSTH) of the firing rate for ‘paired’ excitatory odor responses of 3 pSPNs in DAT^ChR2^ mice plotted for ‘pre’ and ‘post’ phases. The responses of each unit were normalized to 1. (h) Mean PSTH ± standard error of pSPNs with excitatory responses to the paired (left) and non-paired odorant in DAT^ChR2^ mice (paired t-test corrected for multiple comparisons across bins, asterisk indicates significance) (see also SI Fig. 4e-f).

These observations support that neuronal representations evoked by the presentation of a stimulus are modified when the animal learns to assign value to this stimulus. Furthermore, the neuronal responses evoked by rewarded and non-rewarded stimuli become more distinct during learning. Phasic DA is thought to be released in response to reward retrieval and predicted to induce plasticity to population encoding for the rewarded stimuli. The present observations are compatible with such a phenomenon. Yet, in behavioral tasks, it is difficult to isolate the direct effects of DA. First, it cannot be easily disentangled whether changes in stimulus encoding are evoked by direct effects of DA on the striatal network or by systems effects. The latter comprise different states of motivation or, for instance, preparation for reward retrieval that may also be represented in the same network. Secondly, the manipulation of DA release will change learning behavior and, consequently, neuronal coding. Finally, even with a delay window, the actual stimulus response cannot be easily separated from the activity related to reward anticipation or action preparation. We designed therefore a paradigm aiming to isolate and understand pDA induced plasticity in sensory representations in ventral striatal networks.

### Phasic dopamine induces plasticity selective to paired striatal odor responses

To test whether pDA is sufficient to induce plasticity to the population encoding of stimuli, we performed recordings in awake head-fixed mice that had been habituated to the recording setup for at least a week to minimize distress and movement artifacts. Brief 0.5 s bouts of two natural flower odorants were presented to the mice throughout the experiment. Sniffing activity was continuously monitored (Fig. 2a). Recordings were performed with the newly developed tetrode array chronically implanted bilaterally to the OTu. The release of pDA was evoked through fiber optics placed above both OTu in DAT:Cre mice that were injected with a Cre-dependent AAV and expressed ChR2 selectively in dopaminergic midbrain neurons (DAT^CHR2^). Animals with selective expression of GFP in dopaminergic neurons served as a control cohort (DAT^GFP^).

To examine whether pDA evoked plasticity selectively for the paired natural odorant, we designed a protocol consisting of three phases (Fig. 2a). Bouts of two odorants were applied throughout all phases in a pseudo-randomized order. After sampling odorant responses in a ‘pre’ phase, one of the two odorants was paired with a brief burst of optogenetic excitation of dopaminergic terminals in the ventral striatum (12 pulses at 40 Hz) (‘pairing’ phase); then a ‘post’ phase followed to track the further evolution of odorant responses after evoked pDA. We will refer to these two odorants as *paired* and *non-paired* odorants, respectively. Except for the pDA release phase of the paired odorant, all odor presentations were combined with a sham laser light above the animals’ head to account for visual stimulation effects. It is important to note that the recorded mice never experienced the delivery of reward following stimulus presentation, neither prior to the experiment, nor during it. This experimental strategy was designed to dissociate direct DA-induced plasticity from other indirect systems or behavioral modifications occurring during active learning tasks.

We recorded a total of 195 single units in 6 DAT^GFP^ mice and 198 units in 6 DAT^CHR2^ mice. Among them, only units with a baseline firing rate < 5 Hz were considered pSPNs (SI Fig. S3a). To examine pDA induced plasticity, we included units with stable response to odorants throughout the entire ‘pre’ phase (Fig. 2b). Putative SPNs, both in the DAT^CHR2^ and DAT^GFP^ cohort, when responsive, displayed either inhibitory or excitatory responses to the odorants (Fig. 2b). A subset of neurons responded to only one of the two presented odorants and a subset to both, maintaining, in this case, the same response type (either inhibitory or excitatory) for both odorants (Fig. 2b). In both cohorts the timing of the peak response varied among pSPNs in a window ranging from 0 to 1 sec from odor onset (SI Fig. S3b). We quantified the plasticity of the population response by computing the distance from baseline of the population vector during the ‘pre’ and ‘post’ phases. We chose to focus on ‘pre’ and ‘post’ phases to avoid the possible acute effects due to the laser stimulation of the ‘pairing’ phase. In DAT^GFP^ mice, no change was observed from ‘pre’ to ‘post’ for either odor response (Fig. 2c, SI Fig. S3c). In DAT^CHR2^ mice, the deflection from baseline increased selectively in response to the paired odorant, while it remained unchanged for the non-paired odorant (Fig. 2d, SI Fig. S3c). The selective plasticity of the paired odor response was also confirmed when neuronal activity was aligned to the onset of the first inhalation after stimulus presentation (SI Fig. S3d).

To specifically address whether the representation itself of the same odorant changed from ‘pre’ to ‘post’, and not only with respect to baseline, we focused on the response window of each odorant (see Methods) and tested whether the change in odor responses after pDA paring exceeded the trial-by-trial variability within the ‘pre’ phase. To this aim, we tested if the distance between the trial responses of the ‘pre’ and ‘post’ phases was bigger than the trial-by-trial variability within the ‘pre’ phase. Indeed, the representation of the paired odorant changed as result of the pDA paring in DAT^CHR2^ animals only (Fig 2e-f, SI Fig. S3e-f). This effect was robust to randomization of the trial order across sessions (SI Fig. S4a-d) and significantly changed both for the cosine and the Euclidean metric (Fig 3, SI Fig. S3–4). The neuronal representation of the non-paired odorant remained largely unchanged (Fig. 3f, SI Fig. S3f). The effect of pDA stimulation on the paired odorant persisted without further stimulation for the recorded duration of 20 min (‘post’ phase). A further extension of the ‘post’ phase was not possible in our experimental conditions as in sessions longer than 50 min in total, mice became frequently drowsy and neuronal baseline firing rates changed globally (data not shown).

**Figure 3.**
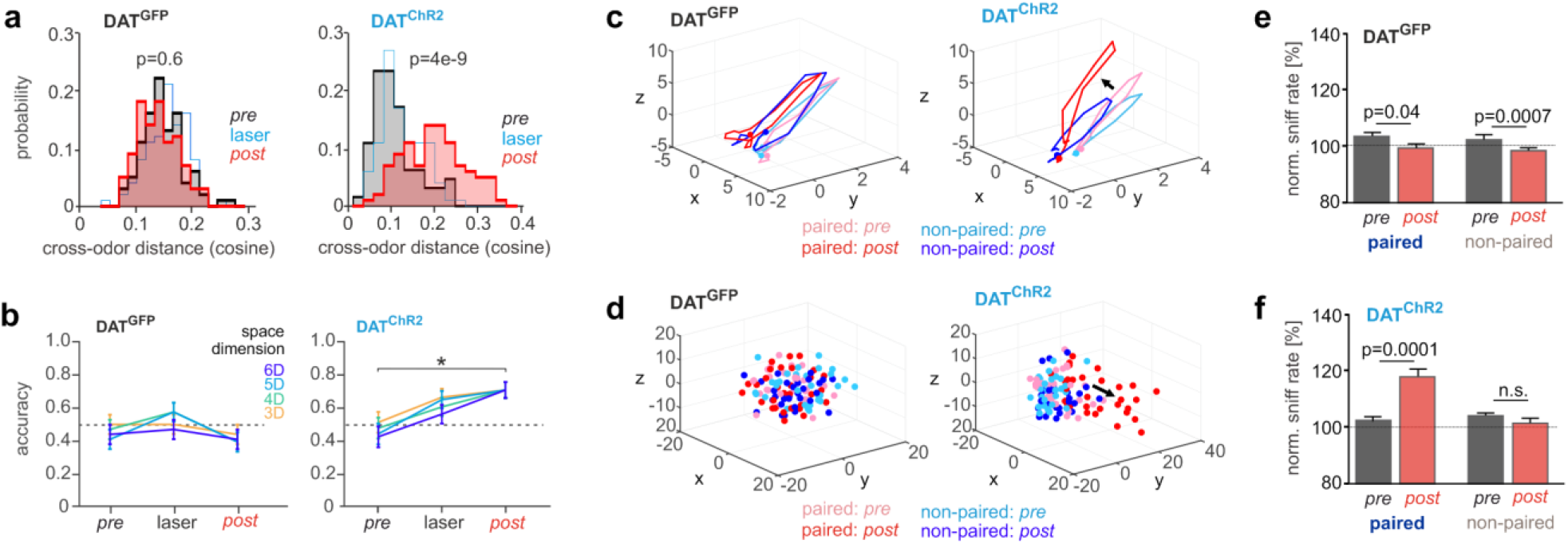
Phasic DA increases the difference between the paired and non-paired odor encoding and improves decoding. (a) To test if the distance between the neural representations of the two odorants increased after pDA paring, we computed the cosine distance between the response vectors of the two odorants during ‘pre’ and ‘post’ phases. After pDA paring, cross-odor distance increased only in DAT^ChR2^ mice. Effect tested with two-way ANOVA (factors: cohort and odor). Interaction effect: F(1,360)=90.6 p=3×10^−19^, post-hoc comparisons (Tukey’s correction) are indicated. (b) We tested if pDA improved odor discriminability by performing a quadratic discriminant analysis between paired and non-paired odor responses. The average accuracy ± standard error of the classifier across the test set is reported. Only in DAT^ChR2^ mice the accuracy of the classifier increased after pDA paring. Multiple state space dimensions are tested to confirm the robustness of the effect. Significance was tested with Fisher’s exact test corrected for multiple comparisons. (c) Averaged trajectories for paired and non-paired odor responses for pSPNs in DAT^ChR2^ and DAT^GFP^ mice. Trajectories, after time embedding expansion, are visualized in 3D through factor analysis (a dot marks the beginning of the trial). Note the selective change of the population vector after pairing of the odorant with pDA. (d) Trial-specific odor responses visualized through multidimensional scaling. Note that after pDA paring, the responses to the paired odorant separate both from the non-paired-responses and from the responses of the paired odorant itself before pairing. (e) Mean sniff rate ± standard error following odor-onset normalized to baseline (computed directly before the odorant application). Data are plotted as mean across DAT^GFP^ animals before and after the pairing protocol (paired t-test). (f) same as (e), but for DAT^ChR2^ mice. In the ‘post’ phase, only the odorant paired with pDA stimulation increases its salience for the animal.

Both sub- and suprathreshold changes in activity to the stimuli contribute to the population dynamics captured above. We tested whether we would observe plasticity in the suprathreshold responses in pSPNs. pSPNs displayed different response types, we split them respectively into dominantly excitatory, inhibitory and non-responsive units for each odorant. Among pSPNs with a significant odor response, we observed a change in firing rate selectively for those with excitatory responses and only for DAT^CHR2^ mice (Fig. 2g-h, SI Fig. S4e-f). After the pairing phase, this subset increased its firing rate in response to the paired odorant, while no change occurred in response to the non-paired odorant. Thus, pDA evoked plasticity in the neuronal representation selectively for the paired sensory objects.

### Phasic dopamine increases the discriminability of striatal odor responses

Above we have shown that the neural representation changes after pDA stimulation only for paired odorants. We were therefore interested in establishing if such plasticity would move the odorant representation closer to the non-paired one, or farther apart. We compared the distance between the population responses to paired and non-paired odorant before and after pairing. In DAT^GFP^ mice, the distance between the two neuronal representations did not change throughout the phases (Fig. 3a, SI Fig. S5). In contrast, in Chr2^DAT^ mice, pairing with pDA increased the distance between the paired and non-paired odor representations (Fig. 3a, SI Fig. S5). To explicitly test whether this effect could enhance discriminability between the two odorants we performed a quadratic discriminant analysis on the population vectors of the paired and non-paired odor responses during the ‘pre’ and ‘post’ phases. The classifier was obtained by first estimating on the training set the mean and covariance of two multidimensional Gaussian distributions, one for each odorant. The class membership of the odor responses of the test data set was assigned on the basis of their Mahalanobis distance from the two Gaussian distributions. In order not to overfit data, we reduced the dimensionality of the population space (see Methods). To make sure that the obtained results did not depend on the precise dimensionality of the reduced space, we repeated the test for all space dimensions for which the number of model parameters was smaller than the sample size. We found that in DAT^CHR2^ mice the accuracy of the classifier significantly improved after pDA pairing (Fig. 3b right). For DAT^GFP^ mice, instead, no change occurred (Fig. 3b left).

The plasticity elicited by pDA paring altered selectively the evolution of the population trajectory of the paired odor response (Fig. 3c). Such deflection had the effect of both increasing the odor response, by increasing its departure from the resting condition (Fig. 3c, cf. Fig. 2d), and setting it apart from the representation of other odorants (Fig. 3d, cf. Fig. 3a).

### Phasic dopamine increases the perceived salience of paired odorants

As we had found that pDA induced plasticity selectively to the coincidently presented stimulus and increased its distinctness from other stimuli, we asked whether the animals may perceive the paired odorant differently than the non-paired odorant. To quantify the perception of the odorant, we recorded the sniffing frequency of the animal and used it as proxy for the perceived salience of the stimuli (38, 39).

We recorded the sniffing frequency in DAT^GFP^ and DAT^ChR2^ mice with an odor port mask. The mask allowed for measurement of the breathing cycle as the system was calibrated to keep the (odorized) air inflow constant relative to the outflow. Throughout the experiment, sniff frequency remained constant (SI Fig. S6a-b). A small increase in response to the odorant presentation habituated over the duration of the recording session in DAT^GFP^ mice (Fig. 3e, SI Fig. S6c). Yet, consistent with the hypothesis that the perceived salience increases through pDA, the sniff frequency increased selectively to the paired odorant after evoked pDA release in DAT^CHR2^ mice (Fig. 3f, SI Fig. S6d). In summary, pDA is sufficient to induce plasticity to make natural stimuli more salient over others, both in their neuronal representations and perception.

## Discussion

The ventral striatum comprises functionally segregated circuits for processing of wanting, liking, drug seeking, aversive and incentive salience behaviors (4, 25, 35). The observations in this study on valence coding of olfactory cues, together with other evidence (34, 35, 40), establish a specialized olfactory stream within the ventral striatum. Phasic DA was sufficient to induce plasticity in the dynamics of the population encoding that matched modifications of the conditioned stimulus representations during reinforcement learning. The underlying modifications in the neuronal representations happened selectively to the odorant which had been paired with pDA release, while the response to the non-paired odorant remained relatively unchanged. Through this direct dopamine induced plasticity, the population representation of the paired odorant stuck out both from baseline network activity and from other odor representations as also observed during reinforcement learning. The stimuli were natural flower odorants composed of many molecules, suggesting that pDA modifies the encoding of complex odor objects as experienced naturally by the animal. Compatible with the observation that striatum primarily encodes stimulus value (25, 26, 41), before pairing with pDA, the striatal network did not distinguish in its population encoding the initially neutral stimuli of different odor identities (Fig. 3b). Pairing with dopamine was therefore sufficient to persistently improve the distinctness in the decoding of the previously paired and non-paired odorants, compatible with demands of neuronal representations of increased salience. The dynamics in the population code capture all changes in the network response including complex supra- and subthreshold changes in single units (18). Among the suprathreshold responses, pDA enhanced the intensity of excitatory stimulus responses; a modification that was also observed during reinforcement learning in pSPNs in this study. This in vivo plasticity is compatible with synaptic plasticity in vitro through DA receptor activation in SPNs (42–44) that increases the firing output to olfactory input stimulation (44). Interestingly, the dopamine-induced plasticity in the reward system employs different network and cellular modifications from the auditory cortex of anesthetized rats, where the cortical area of tone representation increased, but the discharge rate of neurons to stimulus remained unchanged (45).

### Network plasticity and behavior

The DA-induced plasticity provides a mechanism to increase saliency by enhancing population encoding and distinctness of stimuli. Even though our experimental strategy was such that the animal never experienced a physical reward to the stimuli, the enhanced sniffing revealed an increase in the perceived salience. The selective enhancement of perceived salience persisted after the pairing with the evoked phasic dopamine had ceased. Together this indicates that dopamine alone is sufficient to make stimuli more salient both at the cognitive and network encoding level.

These findings may help to understand dopamine actions both in physiological behaviors and psychiatric disorders. For instance, learnt natural behaviors like the formation of social bonds rely on DA release in the ventral striatum (3, 5, 13). Future studies may test whether the observed plasticity may induce the formation of odor engrams of social partners proposed by Walum and Young (3). In psychiatric disorders like addiction, environmental stimulus association mediated by DA-dependent plasticity in ventral striatal synapses, plays a key role in evoking drug seeking and relapse (4–8). DA-induced plasticity in striatal stimulus representations may provide a corresponding network mechanism of cue conditioning, resulting in increased perceived salience of drug-associated triggers.

## Methods

### Animals and husbandry

C57BL/6J (Charles-River) mice were single housed upon implantation of the recording array at a 12 hours’ day-and-night-cycle. DAT:(IRES)Cre mice (B6.SJL-*Slc6a3*^*tm1.1(cre)Bkmn*^/J; RRID: IMSR_JAX:006660) (46) were maintained in a heterozygous C57BL/6J (Charles-River) background. All procedures were approved by the local animal welfare committee and in accordance with NIH guidelines.

### Virus preparation

To generate cell-type specific expression of Channelrhodopsin 2 (ChR2), we injected a Cre-inducible recombinant adeno-associated virus (AAV) vector rAAV_1/2_-double floxed(DIOA)-ChR2:mCherry (47) into heterozygous DAT:Cre mice. For control experiments, DAT:Cre mice were injected with a rAAV_1/2_-DIOA-eGFP virus (47). Cre-inducible recombinant AAV vectors were produced with AAV1/2 coat proteins to a final viral concentration of 10^16^ genome copies/ml. DAT:Cre positive mice 8 weeks of age were anesthetized with isoflurane; and 0.75 µl of purified virus was injected into each hemisphere of the ventral midbrain (location from Bregma: posterior, 3.0 mm; lateral, 0.8 mm; ventral, 4.4 mm). All mice recovered for at least 21 d before undergoing implantation of the recording array. Expression was confirmed in post-hoc histological exam. Selectivity of viral overexpression to TH positive ventral midbrain neurons had been previously determined (Wieland et al., 2014) and was confirmed in the present cohort (not shown), generating mice with ChR2:mCherry expression in dopaminergic neurons (‘ChR2^DAT^ mice’) or control mice (‘GFP^DAT^’).

### Recording array

For the recordings during reversal learning, the OTu was targeted with 16 tetrodes per array and for the DA plasticity experiment each array contained up to 32 tetrodes. Given the depth of the target structures (>5 mm) and curvature of the OTu, we developed an array (SI Fig. S1c) to address these challenges. The design exploits printed circuit boards (PCB) (±10 µm, Würth Electronics) to achieve a precise x-y-placement of the guiding channels for tetrodes (spun from 12.5 µm teflon-coated tungsten wire, California Fine Wire) and optical fibers (Thorlabs FT-200-EMT). As a building scaffold, six identical PCBs were stacked precisely on top of each other (SI Fig. 1c) via 26-gauge steel tubing threaded through four holes placed peripherally. This allows for placement and parallel alignment of multiple recording electrodes (and optical fibers) in the guiding channels formed by the aligned holes. The lowermost PCB in the scaffold serves as a base plate with plugged holes, assuring z-axis alignment. An identical PCB stacked on top of the scaffold and equipped with a soldered-on Molex SlimStack connector (mated height: 1 mm, pitch: 0.4 mm, 70 channels, model n. 502426-7030), thus becoming the electrode interface board (EIB) and eventual implant. For our application, the board was placed above a molding of the rostro-ventral brain surface, so tetrodes and an optical fiber matched the bowl-shaped 3D-curvature of the OTu. The tetrodes were then fixed in place with a drop of liquid acrylic adhesive. The single wires were connected by threading them through 200 µm vias and through-hole to achieve reliable mechanical and electrical contacts with the EIB. To protect them from damage, they were encased in two-component epoxy. After assembly of the recording arrays, tetrode tips were gold-plated to achieve a lower recording impedance (target: 300 kOhm at 1 kHz) with a NanoZ-device (Multichannelsystems). To connect to the Intan RHD2164 head stage during recordings, a custom-built adapter from the Molex SlimStack connector to two 36 Omnetics Nano Strip connectors was used.

### Implantation of recording array

Mice were anesthetized with isoflurane and pre- and post-surgery analgesia was administered. A roughly circular patch of skin above the skull was removed. Local anesthesia was applied to the skull. The lateral and nuchal muscle insertions were left intact. The operative field was then prepared by attaching the margins of the remaining skin to the circumference of the top of the skull with VetBond (3M), thus protecting soft tissue from damage, contamination or necrosis and leaving only the surface of the skull exposed. Holes were drilled in the skull above the regions where tetrodes and fiber optics were inserted and for grounding above cerebellum. The skull was then coated with Super-Bond C&B (Sun Medical) following insertion of a small Neuralynx gold pin as ground connected with the recording array through an insulated copper wire. The tetrode array was slowly lowered into the brain with a motorized 3-axes micromanipulator (Luigs&Neumann). The center of the tetrode bundle was targeted from Bregma to anterior 1.6 mm and lateral 1.3 mm; reaching ventral 4.9 mm from dorsal brain surface. After reaching target depth, dental cement (Kulzer Palladur) was applied at the margins of the recording array so that gravity and capillary force ensured complete filling of the narrow gap between the bottom of the array and the adhesive-coated skull. After the cement hardened animals recovered in their home-cage. The entire surgical procedure took approximately one hour. Animals were normally fit and eating after less than thirty minutes.

### Confirmation of electrode placement

After completion of the experiments, animals were euthanized and perfused with 4% PFA. Explanting the brain, skull, and implant in toto confirmed rostrocaudal and mediolateral placement in the borders of the OT (SI Fig. 1a). Due to tissue shrinkage and low levels of scar formation and microglia activation, we could not reliably detect fiber tracts in histological sections with Nissl stain or immunohistochemistry against glial proteins with the tetrode array (not shown). We therefore followed for the present experiments a different approach: Heads were severed with the implants attached and subsequently soaked in 4% PFA for up to 4 weeks. Due to tissue shrinkage, the exact location of the tetrode tip could not be determined, but revealed preserved electrode tracts in sagittal 150 µm sections (SI Fig. 1b).

### Recording configurations

After recovery, the mice were placed in the head-fixation setup. The first few sessions were brief (5–20 min) and served to habituate the animals to head fixation in the setup. Mice typically remained mostly quiescent after 1–2 sessions of head fixation, after which odor sessions started. For odorant delivery, a custom-built air-dilution olfactometer was used (Shusterman et al., 2011). Odorants were kept in liquid phase (diluted 1:100 in mineral oil) in dark vials and mixed into the nitrogen stream that was further diluted 1:10 into a constant air stream in the olfactometer. The following natural flower odorants were used in the dopamine plasticity experiments: geranium and ylang-ylang (Sigma-Aldrich W250813 and W311936, respect.). These two odorants were selected based on their shared low subjective trigeminal component and because they evoked comparable stable and broad responses throughout the recording sessions. Odorants were delivered in a pseudo-randomized order with a maximum of three consecutive presentations of the same odorant. Odorants were applied for 500 ms with an inter-trial interval of 10 s. Recordings were performed with two Intan 64 channel RHD 2164 miniature amplifier boards connected to a RHD2000 interface board and open-source Intan interface software. Inputs from the laser, olfactometer and sniff sensor were simultaneously recorded with the same interface board, as were reward application and licking activity (via an infra-red beam break sensor positioned in front of the licking spout, Omron Electronics EE-SX3009-P1). Data were sampled at 30 kHz.

### Sniff recording

To monitor the sniff signal, we used a custom-built snout mask that was gently pressed against the snout to generate a cavity in the mask in which pressure fluctuations were continuously measured through a HDI pressure sensor HDIM020GBY8H3 from First Sensor Inc. connected to the analog input of the Intan RHD2000 interface board. The mask design is modified based on the original design of Dmitry Rinberg. The influx of odorized air into the cavity of the mask was calibrated to the outflow through continuously measured vacuum. The animals quickly adapted and tolerated the pressure mask.

The sniff signal was converted and analyzed with custom-written MATLAB scripts using a 1 to 50 Hz band pass filter. For monitoring changes in the sniff frequency, we analyzed the duration of two full sniff cycles before and after odor onset. Averages were formed for each recording the ‘pre’ and ‘post’ phase of each recording session. We did not analyze the amplitudes of the sniffing as it may be confounded for instance by minor snout movements that may result in pressure fluctuations in the cavity.

Trials were aligned to the sniff cycle by finding the first onset of inhalation after odor presentation and then adding this time difference to the odor onset of each trial. For the sparse firing of pSPNs there was no marked effect by sniff alignment. The results in pSPNs for DA induced plasticity were also observed after sniff alignment (SI Fig. S3d). Yet, for the multiphasic and highly active FS neurons, we consistently performed all analysis only after sniff alignment.

### Optogenetic plasticity experiments

The plasticity experiments involving evoked phasic DA release followed a generic design with three phases: Two odorants were applied in a pseudorandomized fashion for 500 ms with an inter-trial-interval of 10 s. In each phase no odorant was consecutively applied more than three times in a row. We initially pseudo-randomly varied the number of trials per phase around 30. In later experiments we consistently used 40 trials in the first phase (the first 10 were omitted for the analysis), 30 trials in the second phase and 30 trials in the third phase. The odor responses before pairing with evoked DA release were recorded in the ‘pre’ phase, followed by a ‘pairing’ phase during which one of the two odorants was paired with a train of 5 ms laser pulses at 40 Hz for 300 ms starting simultaneously with the onset of the odorant application. TTL-driven laser pulses (5 ms duration, 2 to 5 mW at fiber tip) were delivered from 200 µm multimode optical fibers (Thorlabs) coupled to a 25 mW, 473 nm, diode-pumped solid-state laser. To then monitor changes in responses, odorants were continued to be applied in the ‘post’ phase. Usually two sessions were performed in each ChR2^DAT^ or control GFP^DAT^ mouse, and the odorant that was paired with the ‘pairing’ laser, was switched from session to session. A second laser provided ‘sham’ stimulation above the head of the animal with the same intensity and frequency in all other trials, except when the ‘pairing’ laser was on. Additionally, blue light illumination was constantly on in the recording chamber.

### Behavioral conditioning in the go/no-go task

We trained the animals in the head-fix setup described above. Mice received water in their home cage so that their body weight stabilized at 85% of baseline body weight. The training comprised multiple stages and progressed after reaching a performance criterion defined as at least 80% correct responses in 40 consecutive trials. Trials were considered correct if either at a ‘go’-response the reward was retrieved or at a ‘no-go’-response no significant licking was detected during the lick window. In the initial sessions, the animals’ licking behavior was shaped by first presenting them with a drop of water and subsequently letting them obtain more water when they licked at the licking spout (available in a random interval schedule, 0.5-12 s). Stage 1 (‘training’): A single odorant (1.5 s stimulus duration) was presented. Animals could obtain a 5 µl drop of water if they licked at least three times during a window from 0 to 2.5 s after stimulus onset (‘retrieval window’), this was considered a ‘go’-response. The interval between trials was randomly set at 10±2 s in all stages. Stage 2 (‘discrimination’) consisted of two odorants in pseudo-random succession (no more than 3 consecutive trials with the same stimulus). One odorant (1.5s duration) was rewarded as in stage 1 (retrieval window: 0.5 – 2.5 s), while a ‘go’-response for the second odorant was registered as a false alarm and thus incorrect. No punishment was used. Stage 3 (‘reversal learning’) used the same parameters as stage 2, but upon reaching the performance criterion (in the ‘original phase’), the reward contingency of the odors was switched (‘reversal phase’). The data set used in this study consists of one reversal learning session per animal (10 sessions with a total yield of 75 single units) after we accustomed them gradually to a longer lick delay (1.5 – 3 s), keeping odorant delivery concurrent (3 s duration).

### Data pre-processing: Spike detection

To reduce noise and movement artifacts affecting all recording sites, we subtracted the median voltage trace of all channels from each recorded trace. The resulting signal was band pass filtered between 300 and 5000 Hz (4th order Butterworth filter, built-in MATLAB function). A threshold value for spikes was computed as a multiple (7.5x) of the median absolute deviation of the filtered signal (48). Temporally proximal detected peaks over threshold were pruned by height to a minimum distance of 1 ms to avoid multiple detections of the same multiphasic spike. When an event was detected on multiple channels of a tetrode, the timestamp of the highest detected peak was used. Spike waveforms were extracted around −10 to +21 samples around the peak.

### Data pre-processing: Spike sorting

Spike sorting was done with a custom-built graphical user interface in MATLAB, originally developed by A. Koulakov (CSHL). Metrics used for clustering included detected peak height or amplitude (and the respective principal components over channels), and the first three principal components of the waveforms for each respective channel when a spike was predominantly recorded on one channel. The quality of single unit clusters was assessed using the mlib toolbox by Maik Stüttgen (Vs. 6, https://de.mathworks.com/matlabcentral/fileexchange/37339-mlib-toolbox-for-analyzing-spike-data) with particular attention to peak height distribution (fraction of lost spikes due to detection threshold), contamination (fraction of spikes during the refractory period <5ms) and waveform variance.

For analysis of the silicon probe data, spike detection and spike sorting was done using KiloSort (49), followed by manual curation with Phy (https://github.com/kwikteam/phy) using similar parameters to assess unit quality.

### Data pre-processing: Inclusion criteria of units

For further analyses, units were only included if they complied with a set of criteria: Throughout the analyzed part of the recording session units were allowed to only have a maximum change in baseline firing rate from beginning to the end of the session of less than ten percent and intermittent maximum fluctuations of 20%. After exclusion of the first ten trials of odorant application where we frequently observed an initial response habituation, both odor responses had to be stable throughout the ‘pre’ phase. Few units that were responding with short latency to laser pulses in ChR2^DAT^ mice and had features previously described for striatal cholinergic interneurons (50) were excluded from the analyses (not shown).

### Analysis of single unit activity

Single units were divided into two groups: pSPNs and FS. Units with less than 5 Hz baseline firing rate were classified as pSPN (SI Fig. 3a) in agreement with previous molecular tagging experiments (37). To analyze whether odor response plasticity can also be detected at the single unit level (Fig. 2g-h), we divided pSPNs in three groups, neurons with an increase, decrease, or no change in firing rate during the 1 s following odorant application onset. Due to the low baseline firing rate, z-scoring or comparable normalization was not suitable for pSPNs, we therefore considered neurons responsive if the mean firing rate for a 250 ms bin in the 1 s response window changed at least twofold in one of the three phases. After log-transformation, a paired t-test was used to compare ‘pre’ to ‘post’ firing rate. The test was corrected for multiple comparisons (on the multiple bins tested) with the Benjamini–Hochberg correction.

### Population analysis: The population vector

To increase the power of our analysis, considering the relatively small cell count per session due to the numerous selection criteria (see main manuscript), we built spike count population vectors by pooling units from different sessions (20, 51, 52). Since we were interested in either paired or non-paired odor responses, we first divided the trials of each session in paired and non-paired trials. Within these two groups, we combined the population vectors of different sessions 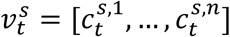, with 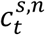 indicating the spike count of unit *n* at time-bin *t* in session *s*, in an across-session population vector *V*_*t*_. The vector *V*_*t*_ was obtained by concatenating 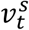 matching the trials according to their trial order, 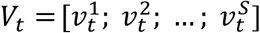. For example, *V*_1_ was obtained by concatenating the first trial of all sessions in which the paired odorant was presented to the animal, *V*_2_ by concatenating the second trial of all sessions with paired odorant, etc. Fig. 1–2, 3a, SI Fig. 2a-c, 3, 5a were produced using population vectors built with such progressive-trial alignment. This trial matching criterion aims to reflect in *V*_*t*_ the temporal progression of the experiment; however, since the units composing *V*_*t*_ were not (all) recorded simultaneously, we wondered if a different order in trial-paring across sessions could lead to different results in our analyses. We then repeated the analyses for 300 different realizations of population vectors 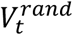 obtained by randomly permuting the trial order in which the session specific population vectors were aligned 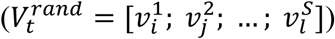. SI Fig. S2c-d, S4, S5b-c were produced using the randomized population vectors. For these latter analyses, the fraction of the 300 performed tests that had a p-value <0.05 is reported in the respective figure legend. Finally, since some sessions had different numbers of trials per phase, we built the population vectors with the minimum number of trials available among sessions. This was done by omitting the last trials of the ‘pairing’ and ‘post’ phases and the initial trials of the ‘pre’ phase.

### Population analysis: Temporal evolution of the odor response

Fig. 1e, 2c-d and SI Fig. 2a-c, and 3c-d show the average deviation of the population vector from baseline following odorant onset. Trials were binned as indicated below. To reduce trial-by-trial variability in neuronal response, we divided the trials in groups of 3 and computed the average population vector of each trial-cluster. In this way, we reduced the number of samples available but we obtained more stable population responses. We did not perform trial grouping for the go/no-go paradigm, given the low number of trials available. For each trial group, we built a baseline population vector by averaging the spike counts of *n*_*bl*_ consecutive baseline bins. We then computed the distance between the baseline vector and the population vector for each bin during the trial. In order to account for changes of the population vector both in direction and in rate, we computed distances with the cosine metric, 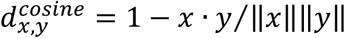, and the Euclidean metric, 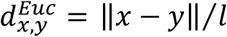 (the Euclidean distance was normalized by the vector length *l*). Finally, we tested the first *n*_*test*_ bins after odor onset for differences in deviation from baseline of the odor responses in the ‘pre’ phase and in the ‘post’ phase with a two-tailed t-test. The test was corrected for multiple comparisons (on the multiple bins tested) with the Benjamini–Hochberg correction. *Parameters:* Behavioral paradigm: *binsize*=0.4 s; *n*_*bl*_ =3 (between −1.8 s to −0.6 s from odor onset); *n*_*test*_=12. Passive paradigm: *binsize*=0.25 s; *n*_*bl*_=4 (between −1.75 s to −1 s from odor onset); *n*_*test*_=7.

### Population analysis: Change in odor encoding after pDA paring

As for the previous analysis, trials were binned in time and averaged in groups of 3. To investigate the effect of pDA on odor responses, we built an odor response population vector by averaging spike counts across the response bins (from 0.6 s to 1.8 s from odor onset in the behavioral paradigm and from 0.5 s to 1.25 s in the passive paradigm). We then computed the distance (both cosine and Euclidean) between all population vectors of each trial-cluster of the ‘post’ phase and ‘pre’ phase and between all population vectors of the ‘pre’ phase within each other. In this way, we could test if the distance between the odor responses in the two phases was bigger than the original trial-by-trial variability within. We finally tested for interaction between cohorts (DAT^ChR2^, DAT^GFP^), odorant type (paired, non-paired) and phase (‘pre’, ’post’) through a three-way ANOVA (independent factors). Post-hoc comparisons were performed with Tukey’s test correction for multiple comparisons (factors: mice group (levels: DAT^ChR2^, DAT^GFP^), phase (levels: ‘pre’, ’post’) and odor (levels: paired, non-paired)). Fig. 2e-f, SI Fig. S3e-f, S4.

### Population analysis: Change in distance between odorant encodings after pDA paring

To test whether the distance between the neural representations of the two odorants increased after DA paring, we computed the distance between the response population vectors (as described above) to the two odorants both during the ‘pre’ phase and ‘post’ phase. Distances were computed with both cosine and Euclidean metrics. Significance was tested with a two-tailed t-test in the behavioral paradigm (where no control group was present, Fig. 1g, SI Fig. S2d-f) and with a two-way ANOVA with post-hoc Tukey’s test correction in the passive paradigm (factors: mice group (levels: DAT^ChR2^, DAT^GFP^) and phase (levels: ‘pre’, ’post’)) in Fig. 3a and SI Fig. 5.

### Population analysis: Odor response classifier

We tested if pDA improved odor discriminability by measuring the accuracy of a classifier in discriminating the odor responses of the paired and non-paired odorants in the ‘pre’ phase and in the ‘post’ phase (Fig. 5b). Trials were divided into four groups based on the odorant (paired vs. non-paired) and of the session phase (‘pre’ vs. ‘post’). Both in the ‘pre’ and ‘post’ phase we performed a quadratic discriminant analysis (QDA) to classify paired and non-paired responses. The first step of QDA is to estimate the parameters of, in this case, two multidimensional Gaussians representing the data of the training set. The number of parameters *df*_*c*_ per Gaussian component scales quadratically with the dimension *D* of the vector space (specifically *df*_*c*_ = *D*(*D* − 1)/2 + 2*D* + 1, yielding in our case *df*_*total*_ = 2 ∗ *df*_*c*_ − 1). Hence, to avoid overfitting, we first reduced the population vector space dimensionality through principal component analysis performed on the four odor response groups simultaneously (making sure that odor representations in the ‘pre’ and ‘post’ phases shared the same space). The classifier was tested with a leave-on-out cross validation procedure, with total accuracy given as *a* = # *correct*/# *tested*. To make sure that the results obtained did not depend on the specific dimensionality of the population space, we repeated this analysis for all dimensions from 3 to 6 (for dim>6, the number of the model free parameters started to exceed the number of data points). Finally, we used Fisher’s exact test to assess whether the number of odor responses correctly assigned by the classifier in the ‘pre’ and ‘post’ phases differed. Significance was corrected for multiple comparisons (Benjamini–Hochberg correction) across the four tested dimensions.

### Population analysis: Visualization of population trajectories

Trials were divided according to the paired and non-paired odorant, and according to the phase of the session: ‘pre’ and ‘post’. Within each of these four groups we averaged the population activity across all trials obtaining four multi-variate time series spanning the trial duration. To better unfold neural dynamics (53, 54), we then applied delay embedding and expanded the state space of the neural trajectories adding, for each unit, *m* delayed coordinates (*m*=3; delay=1 bin). Finally, to visualize the trajectories, we applied factor analysis (FA) and reduced the dimensionality of the space to 3 (Fig. 3c). FA allows describing the observed variables in terms of a reduced number of independent latent factors. To improve visualization, the trajectories were rotated within the axes.

### Population analysis: Visualization of odor responses

Odor response population vectors were defined as explained in ‘*Change in odor encoding after pDA paring*’ and divided in four groups according to odor type and session phase. For visualization purposes, we reduce the vector space dimensionality to 3 (Fig. 3d). To better appreciate within and between group variability we used multidimensional scaling (MDS) as dimensionality reduction method. MDS is a non-linear dimensionality reduction method designed to preserve in the reduced space the pairwise distances between points of the original space. To improve visualization we rotated the position of the points within the axes.

## Author Contributions

Conceptualization, W.K., L.O., E.R., M.S.; Methodology, W.K., L.O., E.R., M.S., and R.S.; Investigation, D.D., D.W., W.K., F.H., C.L., L.O., N.R., E.R, M.S., and S.W.; Writing W.K., L.O., E.R. and M.S.; Funding Acquisition W.K., D.D.; Resources, W.K., L.O., E.R., M.S., R.S.; Supervision, W.K. and E.R.

## Acknowledgements

We thank Dmitry Rinberg for sharing the original design of the odor port mask, Patrick Jendritza and Carla Filosa for their help and Andreas Meyer-Lindenberg for his support. The work was supported by DFG grant CRC 1134 project C04, DFG SPP1665 KE1661/2-2, BMBF grant n. 01GQ1708 and the H. and Ch. Schaller Foundation to W.K and DFG grant CRC 1134 project D01 and DGF DU354/8-2 to D.D.

---

**SI Figure S1.**
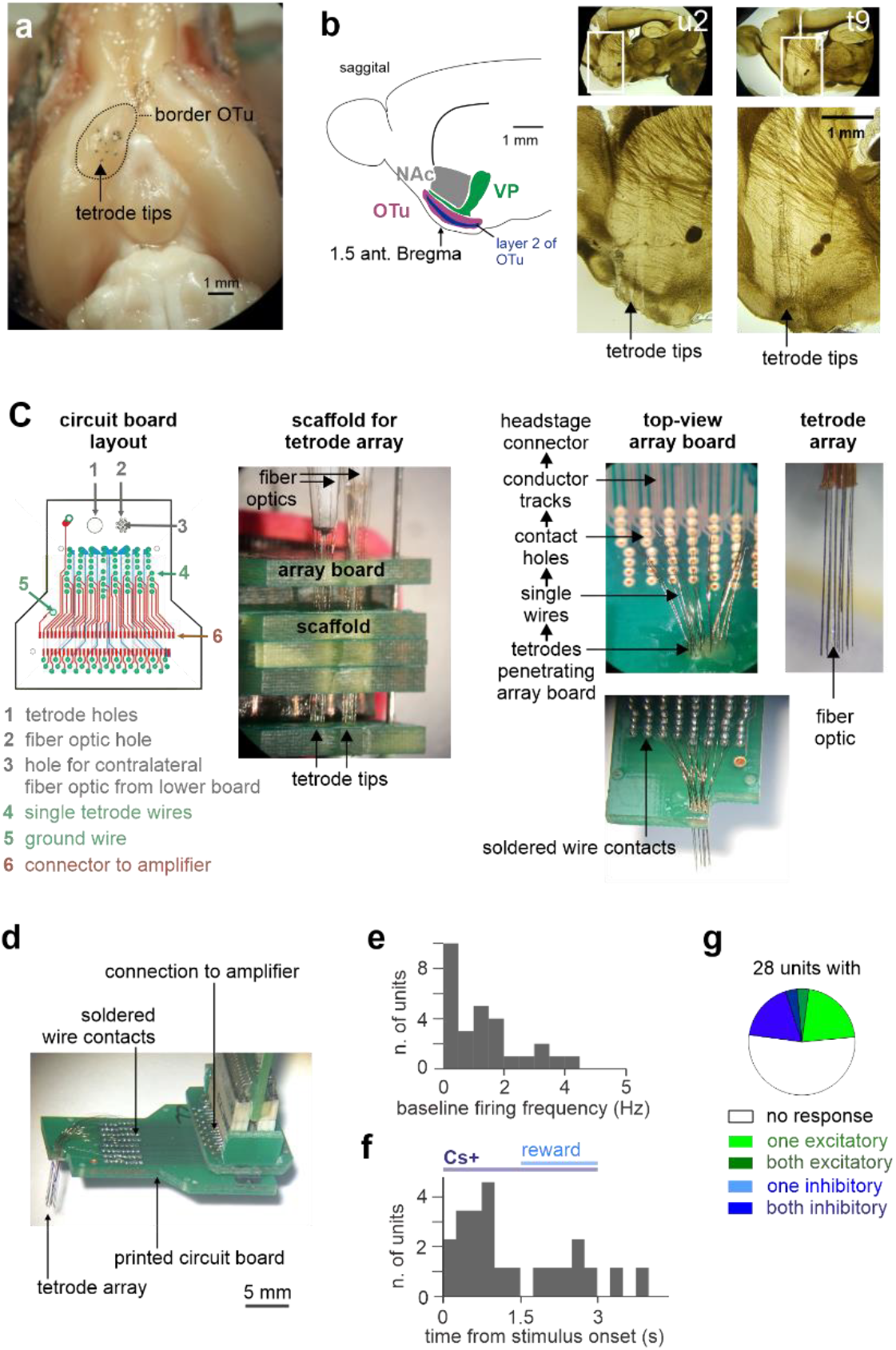
Value assignment to sensory stimuli in the ventral striatum during reversal learning. (a) Histological confirmation of the placement of the tetrodes in the OTu in a ventral forebrain view. (b) Scheme with the anatomical relation of ventral striatal brain regions in sagittal view (left). Histological confirmation of the placement of the tetrodes in the OTu in sagittal sections from two mice #u2 and #t9 (right). (c) Illustration of the building of the tetrode array. Left: Exemplary layout of the printed circuit board. Center left: A scaffold was used to arrange tetrodes in parallel. Center right: Single wires of tetrodes were connected to the board (upper image) and fixed by soldering (lower image). Right: Example of the tetrode array with a fiber optic in the center. (d) The assembled connected to the breakout board of the head stage connector. (e) Baseline firing rates of the pSPN units in the sample (n=28). (f) Time from odorant onset to the peak z-score of the response for pSPN units in the sample (250 ms bins). (g) The fraction of response types to the stimuli is plotted for pSPNs in the OTu.

**SI Figure S2.**
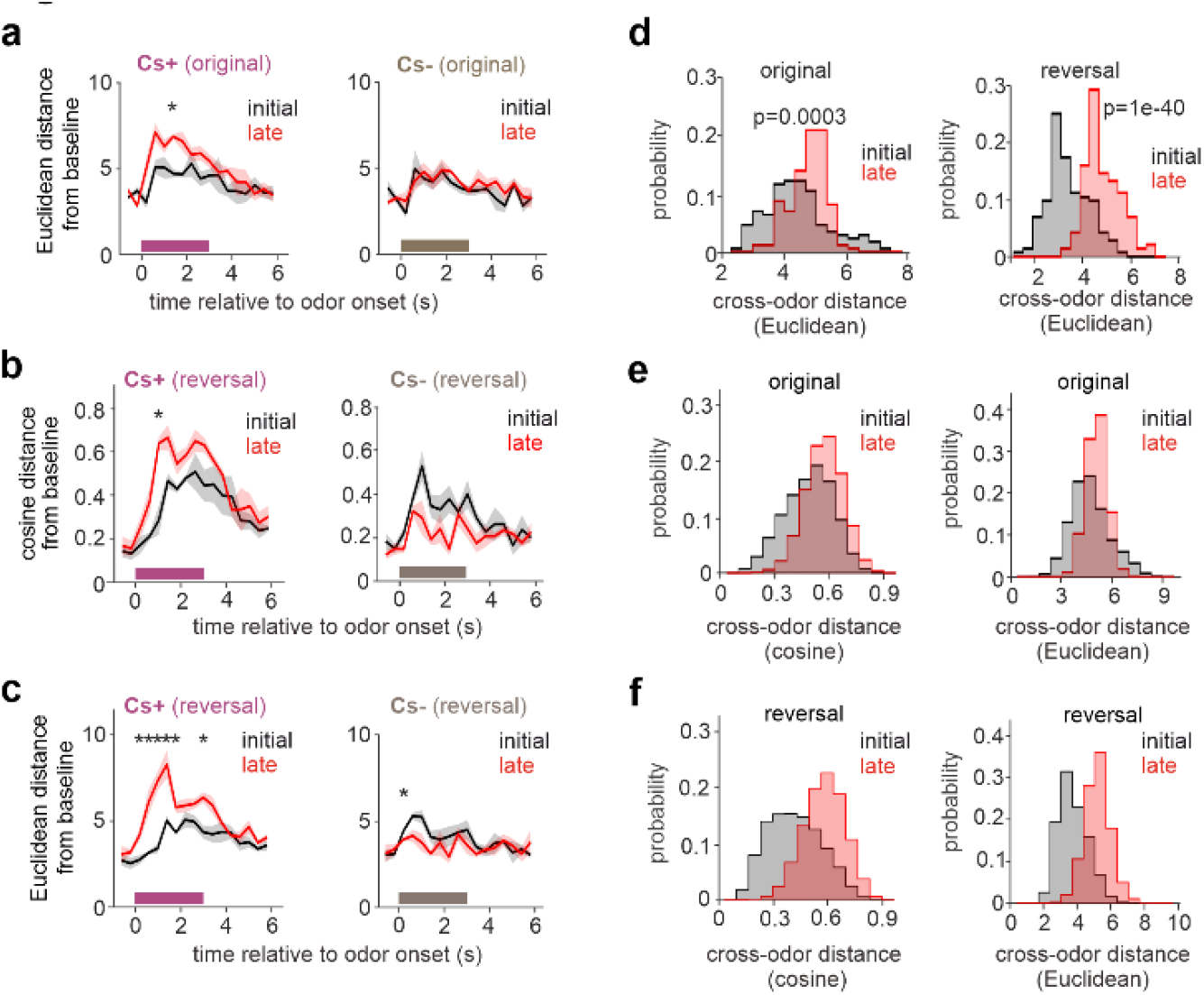
Stimulus related network representations are modified during reversal learning. (a) Mean temporal evolution ± standard error of the Euclidian distance of the population vector from baseline for the rewarded and non-rewarded odorant for the ‘initial’ (black) and ‘late’ (red) phase of the session. Significant changes between the ‘initial’ and ‘late’ phase were assessed with a two-tailed t-test corrected for multiple comparisons across bins. During learning the population vector for Cs+, but not for Cs-, significantly increased its distance from baseline. (b) same as Fig. 1e, but computed on the trials following the reversal of the odorant-reward pairing. (c) same as SI Fig. 2a, but computed on the trials following the reversal of the odorant-reward pairing. (d) Distribution of distances between the trial-specific neuronal representations of Cs+ and Cs-within all trials of the ‘initial’ phase (black) and all trials of the ‘late’ phase (red). Distances were computed both with a, Euclidian metrics. Significance was assessed with a two-tailed t-test. During learning the population vectors relative Cs+ and Cs-diverged. Left: original phase. Right: reversal phase. (e-f) To exclude that the results obtained in Fig. 1g-h depend on a specific trial-alignment in the construction of the population vectors from multiple sessions, we repeated the analyses 300 times on population vectors obtained by randomly permuting the order of trial paring across animals. In 100% of the 300 repetitions the distance between the two odor responses (computed either with the cosine or with the Euclidean metric) increased with learning. The effect was present both during the (e) original and (f) reversal phase.

**SI Figure S3.**
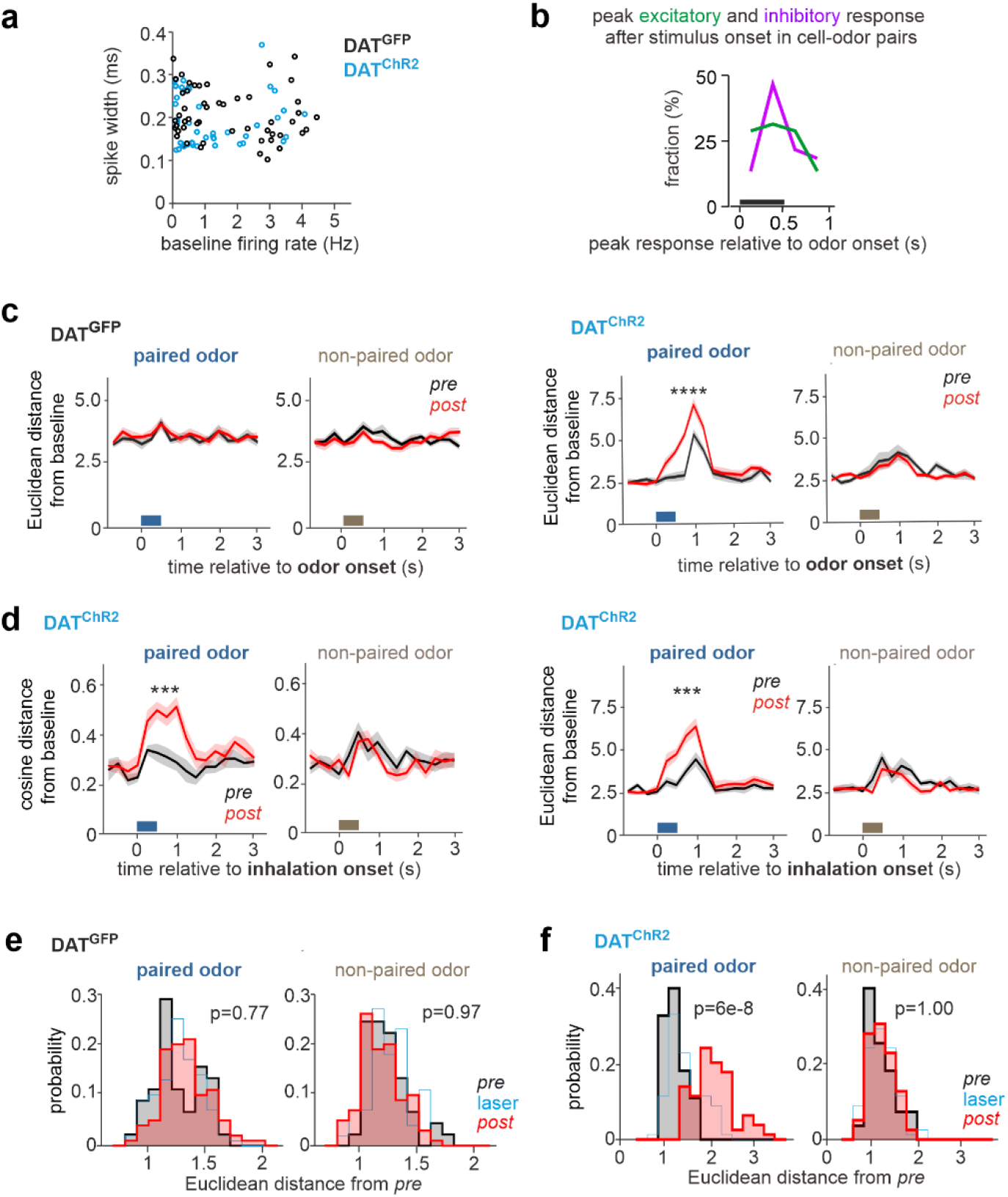
Phasic DA modifies the striatal population code selectively of the paired odorant. (a) The baseline firing rate was plotted against the width at half maximum of the median spike waveform of pSPNs in our sample from DAT^ChR2^ (blue) and DAT^GFP^ (black) mice. (b) For the ‘paired’ odorant in DAT^ChR2^ mice, the time to the peak change in firing rate during the odor response was plotted for all excitatory and inhibitory responses of Fig. 2b (250 ms bins). (c) Same as Fig. 2c-d but computed with the Euclidean metric. (d) Same as (c, right) and Fig. 2d, but computed for neuronal activity aligned to the first inhalation after stimulus onset. (e-f) Same as Fig. 2e-f respectively but computed with the Euclidean metric. Significance established with a three-way ANOVA (factors: cohort, phase, and odor). Interaction effect: F(1,498)=40.3; p=5×10^−10^. Post-hoc tests (Tukey’s correction) are reported on the plots.

**SI Figure S4.**
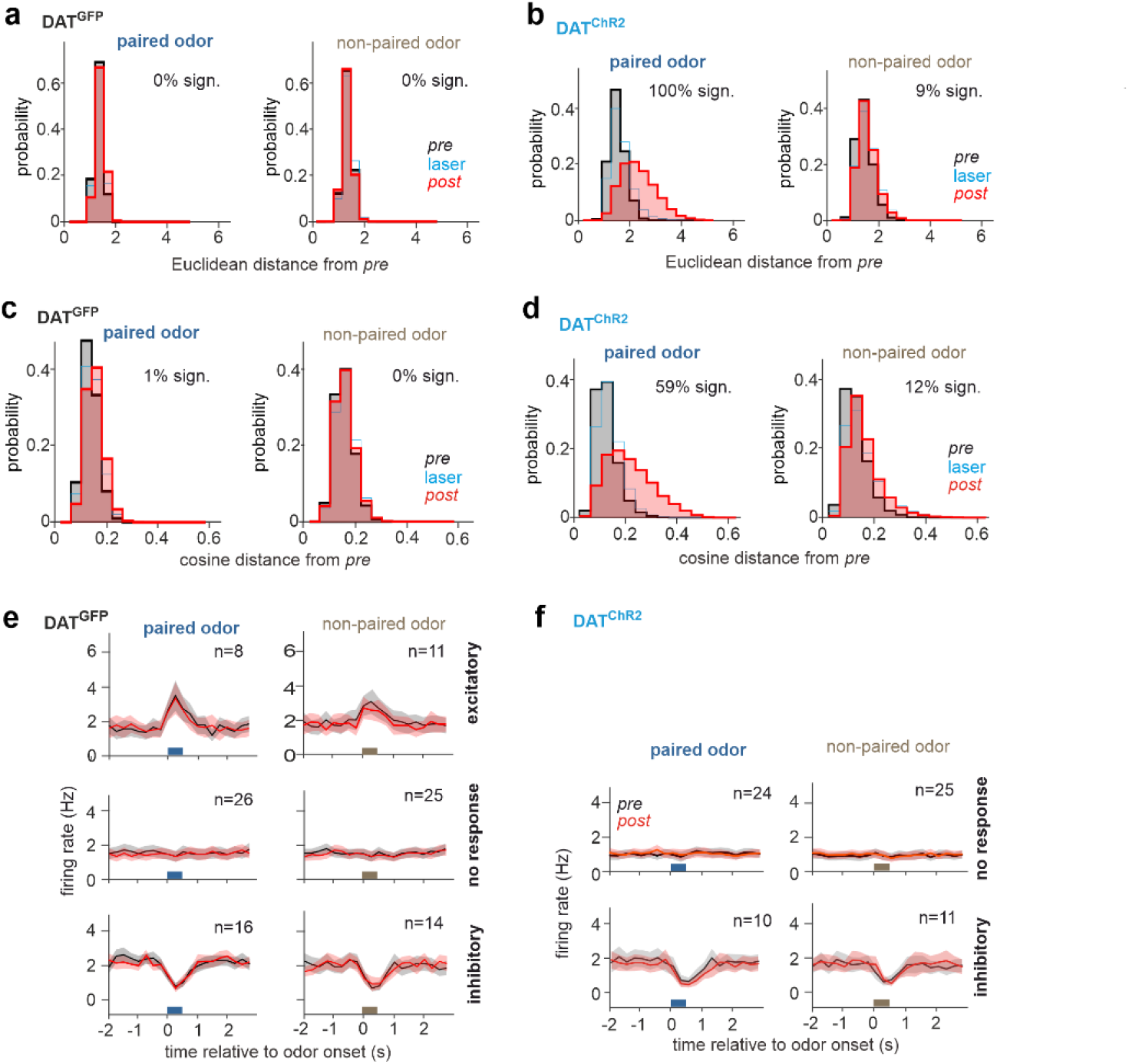
Phasic DA modifies the striatal population code selectively of the paired odorant. (a-b) For the data in Fig. 2e-f, the analyses were repeated 300 times with random permutation, at each repetition, of the order of trial paring across sessions used to build the population vectors with Euclidean metric. For each repetition we performed a three-way ANOVA (factors: cohort, phase, and odor). Of the 300 repetitions the interaction effect between the three factors was found significant 100% of the times in DAT^ChR2^ mice. The fraction of significant post-hoc comparisons (Tukey’s correction) is indicated. (c-d) Same as (a-b) respectively, but with distances computed using the cosine metric. Of the 300 repetitions the interaction effect between the three factors was found significant 59% of the times in DAT^ChR2^ mice. The fraction of significant post-hoc comparisons (Tukey’s correction) is indicated. (e) Mean PSTH ± standard error of pSPNs with excitatory (top), no (middle), and inhibitory responses (bottom) to the paired (left) and non-paired odorant in DAT^GFP^ mice (paired t-test corrected for multiple comparisons across bins). (f) same as (e) for DAT^ChR2^ mice (paired t-test corrected for multiple comparisons across bins, asterisk indicates significance). The excitatory responses are shown in Fig. 2h.

**SI Figure S5.**
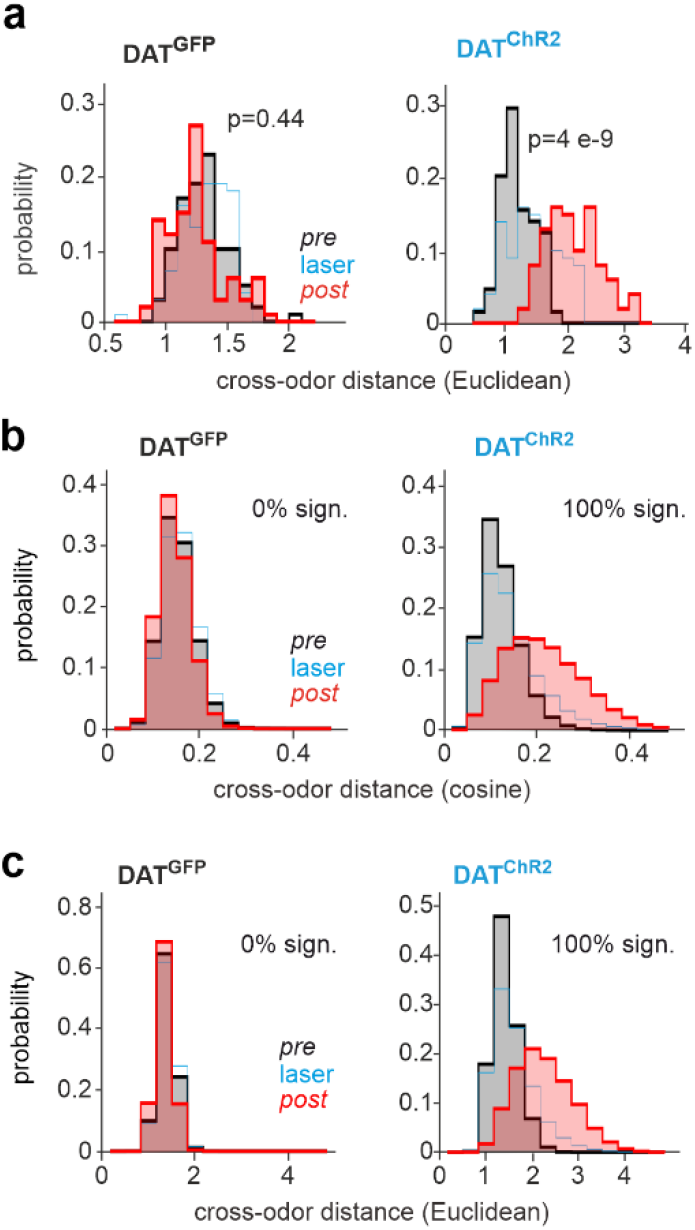
Phasic DA increases the difference between paired and non-paired odor represtations. (a) Same as Fig. 3a but with the use of the Euclidean metric. Statistical test: two-way ANOVA (factors: cohort and odor). Interaction effect: F(1,360)=227.5 p=4×10^−40^, post-hoc comparisons (Tukey’s correction) reported on the plots. (b-c) The analyses performed respectively for (a) and Fig. 3a were repeated 300 times with random permutations, at each repetition, of the order of trial paring across-sessions used to build the population vectors. In DAT^ChR2^ mice, the percentage of tests with significant interaction effect was 100% both for the cosine (b) and the Euclidian metric (c). Fraction of significant post-hoc comparisons (Tukey’s correction) indicated on the figure.

**SI Figure S6.**
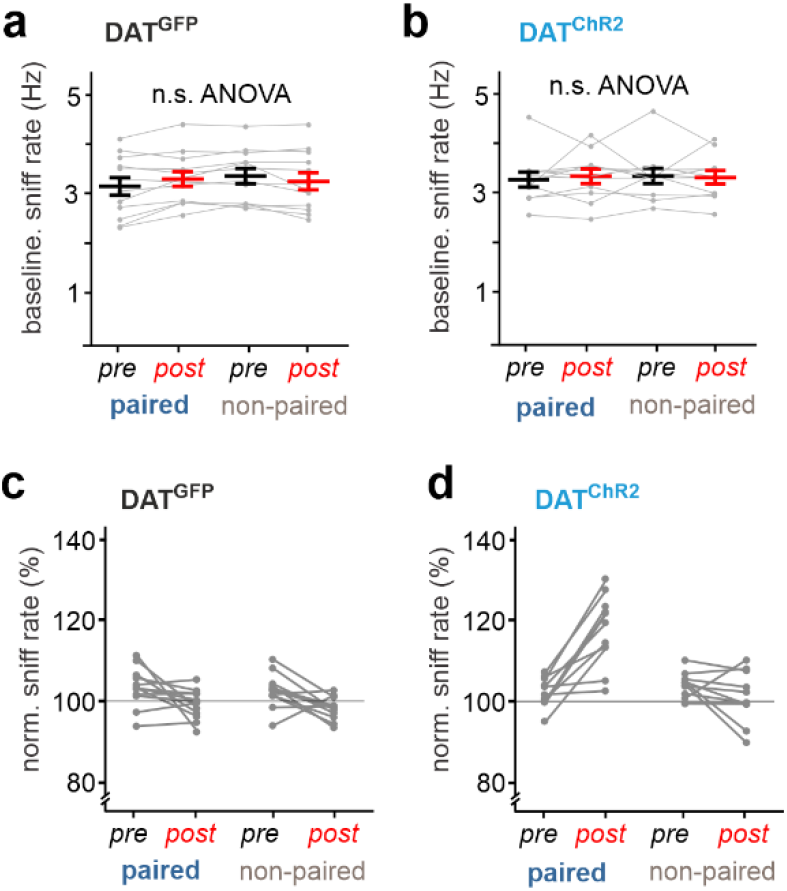
Phasic DA increases persistently the perceived salience of the paired odorant. (a) Mean sniff rate ± standard error directly before the odorant application (baseline rate). Data are plotted as mean across DAT^GFP^ animals before and after the pairing protocol (one-way ANOVA F(3,44)=0.28; p=0.84). (b) Same as (a), but for DAT^ChR2^ mice (one-way ANOVA F(3,40)=0.05; p=0.98). (c-d) The sniff rate following odor-onset normalized to baseline the sniff rate of Fig. 4b-c were respectively plotted separately for each session.

## References

1. Klucharev V, Hytönen K, Rijpkema M, Smidts A, Fernández G (2009) Reinforcement Learning Signal Predicts Social Conformity. Neuron 61(1):140–151.

2. Shamay-Tsoory SG, Abu-Akel A (2016) The Social Salience Hypothesis of Oxytocin. Biol Psychiatry 79(3):194–202.

3. Walum H, Young LJ (2018) The neural mechanisms and circuitry of the pair bond. Nat Rev Neurosci 19(11):643–654.

4. Hyman SE, Malenka RC, Nestler EJ (2006) Neural mechanisms of addiction: the role of reward-related learning and memory. Annu Rev Neurosci 29:565–598.

5. Nestler EJ (2001) Molecular basis of long-term plasticity underlying addiction. Nat Rev Neurosci 2:119.

6. Shalev U, Grimm JW, Shaham Y (2002) Neurobiology of relapse to heroin and cocaine seeking: a review. Pharmacol Rev 54(1):1–42.

7. Zilverstand A, Huang AS, Alia-Klein N, Goldstein RZ (2018) Neuroimaging Impaired Response Inhibition and Salience Attribution in Human Drug Addiction: A Systematic Review. Neuron 98(5):886–903.

8. Graybiel AM, Moratalla R, Robertson HA (1990) Amphetamine and cocaine induce drug-specific activation of the c-fos gene in striosome-matrix compartments and limbic subdivisions of the striatum. Proc Natl Acad Sci U S A 87(17):6912–6916.

9. Kapur S (2003) Psychosis as a state of aberrant salience: a framework linking biology, phenomenology, and pharmacology in schizophrenia. Am J Psychiatry 160(1):13–23.

10. Roiser JP, Howes OD, Chaddock CA, Joyce EM, McGuire P (2013) Neural and Behavioral Correlates of Aberrant Salience in Individuals at Risk for Psychosis. Schizophr Bull 39(6):1328–1336.

11. Redgrave P, Prescott TJ, Gurney K (1999) Is the short-latency dopamine response too short to signal reward error? Trends Neurosci 22(4):146–151.

12. Schultz W (2002) Getting formal with dopamine and reward. Neuron 36(2):241–263.

13. Ungless MA (2004) Dopamine: the salient issue. Trends Neurosci 27(12):702–706.

14. Watabe-Uchida M, Eshel N, Uchida N (2017) Neural Circuitry of Reward Prediction Error. Annu Rev Neurosci 40:373–394.

15. Wise RA (2004) Dopamine, learning and motivation. Nat Rev Neurosci 5(6):483.

16. Rolls ET (2000) On the brain and emotion. Behav Brain Sci 23(2):219–228.

17. Yeshurun Y, Sobel N (2010) An odor is not worth a thousand words: from multidimensional odors to unidimensional odor objects. Annu Rev Psychol 61:219–241, C1-5.

18. Averbeck BB, Latham PE, Pouget A (2006) Neural correlations, population coding and computation. Nat Rev Neurosci 7(5):358.

19. Bakhurin KI, Mac V, Golshani P, Masmanidis SC (2016) Temporal correlations among functionally specialized striatal neural ensembles in reward-conditioned mice. J Neurophysiol 115(3):1521–1532.

20. Laurent G, et al. (2001) Odor encoding as an active, dynamical process: experiments, computation, and theory. Annu Rev Neurosci 24:263–297.

21. Gottfried JA (2010) Central mechanisms of odour object perception. Nat Rev Neurosci 11(9):628–641.

22. Howard JD, Plailly J, Grueschow M, Haynes J-D, Gottfried JA (2009) Odor quality coding and categorization in human posterior piriform cortex. Nat Neurosci 12(7):932–938.

23. Kadohisa M, Wilson DA (2006) Separate encoding of identity and similarity of complex familiar odors in piriform cortex. Proc Natl Acad Sci U S A 103(41):15206–15211.

24. Stettler DD, Axel R (2009) Representations of Odor in the Piriform Cortex. Neuron 63(6):854–864.

25. Berridge KC, Robinson TE (2003) Parsing reward. Trends Neurosci 26(9):507–513.

26. O’Doherty J, et al. (2004) Dissociable roles of ventral and dorsal striatum in instrumental conditioning. Science 304(5669):452–454.

27. Pagnoni G, Zink CF, Montague PR, Berns GS (2002) Activity in human ventral striatum locked to errors of reward prediction. Nat Neurosci 5(2):97.

28. Schultz W (2000) Multiple reward signals in the brain. Nat Rev Neurosci 1(3):199.

29. Radua J, et al. (2015) Ventral Striatal Activation During Reward Processing in Psychosis: A Neurofunctional Meta-Analysis. JAMA Psychiatry 72(12):1243–1251.

30. Dayan P, Niv Y (2008) Reinforcement learning: the good, the bad and the ugly. Curr Opin Neurobiol 18(2):185–196.

31. Montague PR, Hyman SE, Cohen JD (2004) Computational roles for dopamine in behavioural control. Nature 431(7010):760–767.

32. Tsai H-C, et al. (2009) Phasic firing in dopaminergic neurons is sufficient for behavioral conditioning. Science 324(5930):1080–1084.

33. Vetere G, et al. (2019) Memory formation in the absence of experience. Nat Neurosci 22(6):933–940.

34. Zhang Z, et al. (2017) Activation of the dopaminergic pathway from VTA to the medial olfactory tubercle generates odor-preference and reward. eLife 6. doi:10.7554/eLife.25423.

35. Ikemoto S (2007) Dopamine reward circuitry: Two projection systems from the ventral midbrain to the nucleus accumbens–olfactory tubercle complex. Brain Res Rev 56(1):27–78.

36. Zelano C, et al. (2005) Attentional modulation in human primary olfactory cortex. Nat Neurosci 8(1):114–120.

37. Kravitz AV, Owen SF, Kreitzer AC (2013) Optogenetic identification of striatal projection neuron subtypes during in vivo recordings. Brain Res 1511:21–32.

38. Mainland J, Sobel N (2006) The sniff is part of the olfactory percept. Chem Senses 31(2):181–196.

39. Wesson DW, Donahou TN, Johnson MO, Wachowiak M (2008) Sniffing behavior of mice during performance in odor-guided tasks. Chem Senses 33(7):581–596.

40. Gadziola MA, Wesson DW (2016) The Neural Representation of Goal-Directed Actions and Outcomes in the Ventral Striatum’s Olfactory Tubercle. J Neurosci Off J Soc Neurosci 36(2):548–560.

41. van der Meer MAA, Redish AD (2011) Ventral striatum: a critical look at models of learning and evaluation. Curr Opin Neurobiol 21(3):387–392.

42. Reynolds JNJ, Wickens JR (2002) Dopamine-dependent plasticity of corticostriatal synapses. Neural Netw Off J Int Neural Netw Soc 15(4-6):507–521.

43. Shen W, Flajolet M, Greengard P, Surmeier DJ (2008) Dichotomous dopaminergic control of striatal synaptic plasticity. Science 321(5890):848–851.

44. Wieland S, et al. (2015) Phasic Dopamine Modifies Sensory-Driven Output of Striatal Neurons through Synaptic Plasticity. J Neurosci Off J Soc Neurosci 35(27):9946–9956.

45. Bao S, Chan VT, Merzenich MM (2001) Cortical remodelling induced by activity of ventral tegmental dopamine neurons. Nature 412(6842):79.

46. Bäckman CM, et al. (2006) Characterization of a mouse strain expressing Cre recombinase from the 3’ untranslated region of the dopamine transporter locus. Genes N Y N 2000 44(8):383–390.

47. Cardin JA, et al. (2009) Driving fast-spiking cells induces gamma rhythm and controls sensory responses. Nature 459(7247):663–667.

48. Quiroga RQ, Nadasdy Z, Ben-Shaul Y (2004) Unsupervised spike detection and sorting with wavelets and superparamagnetic clustering. Neural Comput 16(8):1661–1687.

49. Pachitariu M, Steinmetz N, Kadir S, Carandini M, D HK (2016) Kilosort: realtime spike-sorting for extracellular electrophysiology with hundreds of channels. bioRxiv:061481.

50. Atallah HE, McCool AD, Howe MW, Graybiel AM (2014) Neurons in the ventral striatum exhibit cell-type-specific representations of outcome during learning. Neuron 82(5):1145–1156.

51. Miura K, Mainen ZF, Uchida N (2012) Odor representations in olfactory cortex: distributed rate coding and decorrelated population activity. Neuron 74(6):1087–1098.

52. Stopfer M, Jayaraman V, Laurent G (2003) Intensity versus identity coding in an olfactory system. Neuron 39(6):991–1004.

53. Balaguer-Ballester E, Lapish CC, Seamans JK, Durstewitz D (2011) Attracting dynamics of frontal cortex ensembles during memory-guided decision-making. PLoS Comput Biol 7(5):e1002057.

54. Takens F (1981) Detecting strange attractors in turbulence. Lect Notes Math Berl Springer Verl 898:366.

